# A Strategy to Quantify Myofibroblast Activation on a Continuous Spectrum

**DOI:** 10.1101/2022.03.09.483659

**Authors:** Alexander Hillsley, Matthew Santoso, Sean M. Engels, Lydia M. Contreras, Adrianne M. Rosales

## Abstract

Myofibroblasts are a highly secretory and contractile phenotype most commonly identified by the de novo expression and assembly of alpha-smooth muscle actin stress fibers. Traditionally, this activation process has been thought of as a binary process, with cells being labeled as “activated” or “quiescent (non-activated)”. More recently, this view has been expanded to consider activation on a continuous spectrum. However, there is no established method to quantify a cell’s position on this spectrum, and as a result, the binary labeling system is still widely used. While transcriptomic analyses provide a continuous measure of myofibroblast markers, a faster and more facile screening method is needed. To this end, we utilized optical microscopy and machine learning methods to quantify myofibroblast activation on a spectrum. We first measured size and shape features of over 1,000 individual cardiac fibroblasts and found that these features provide enough information to predict activation state, on the binary scale, with 94% accuracy as compared to manual classification. We next performed dimensionality reduction techniques on these features to create a continuous scale of activation. Importantly, this new classification system captures a range of fibroblast activation states, but still possesses inherent bias due to choice of morphological features. Thus, we next used self-supervised machine learning to create a second continuous labeling system free from biases associated with the manually measured features. Lastly, we compared our findings for mechanically activated cardiac fibroblasts to a distribution of cell phenotypes generated from transcriptomic data using single-cell RNA sequencing. Altogether, these results demonstrate a continuous spectrum of activation from fibroblast to myofibroblast and provide a strategy to quantify a cell’s position on that spectrum.

## 1. Introduction

The fibroblast-to-myofibroblast transition is a key step in biological processes such as wound healing and the development of fibrotic disease. A range of chemical and mechanical stimuli may initiate this transition, including the inflammatory cytokine TGF-β1^1^ and increased extracellular matrix stiffness^2–4^. Although myofibroblasts can arise from a variety of cell types after an initiating injury, a key source is activation from resident fibroblasts^5^. Upon activation, fibroblasts become increasingly secretory and contractile, eventually adopting the myofibroblast phenotype.

Traditionally, myofibroblasts have been identified by the *de novo* assembly of alpha-smooth muscle actin (α-SMA) stress fibers, most commonly visualized through immunostaining. Typically, this method is used to classify cells on a binary scale, i.e., either “fibroblast” or “myofibroblast.” More recently, however, there is increasing recognition that this binary system is not able to capture the complexities of the full range of transition between these two cell phenotypes. One approach has been to consider activation as a spectrum, including identification of an intermediate phenotype labeled a “proto-myofibroblast,” characterized by diffuse α-SMA expression and α-SMA negative stress fibers.”^6,7^ This spectrum has also been demonstrated through single cell force profiling^8^; however, to date, no system has been developed to quantify the position of individual cells along this spectrum, and as a result, the binary classification system is still widely used. A continuous, rather than binary, classification system would be better able to capture small changes in cell behavior that would be overshadowed by a binary system. For example, a stimulus that causes a partial activation of fibroblasts to a phenotype similar to the “proto-myofibroblast” would not be recognized by the binary system because there was not a significant increase in α-SMA stress fiber positive cells. However, a continuous classification system would be able to capture this more subtle change in phenotype, potentially lending mechanistic insight to fibrotic disease progression.

In addition to the aforementioned concerns, recent work has suggested that the appearance of α-SMA stress fibers is not the only, or even the best, marker of myofibroblast activation^9^. Other markers include expression of myofibroblast specific genes such as collagen type 1^10^, paxillin^11^, or periostin^12^. The assembly of super-mature focal adhesions^7^ has also been associated with myofibroblast activation, as well as the deposition of ED-A fibronectin^13^. Another correlating factor is overall cell morphology, though this is rarely the primary driver of classification^14–16^. For example, it is widely accepted that activated myofibroblasts are significantly larger than non-activated fibroblasts^17–19^, usually have a higher aspect ratio due to cell spreading and possess a less rounded nucleus^20^. However, these metrics have not been rigorously incorporated into models to identify the degree of myofibroblast activation.

Machine learning algorithms offer a way to classify subtle differences across a range of cell phenotypes. Relatively simple algorithms such as decision trees, k-nearest neighbor (kNN), and support vector machines (SVM) have been used to classify blasts in the blood of leukemia patients^21^ and to recognize different types of white blood cells^22^. More complex algorithms such as convolutional neural networks (CNNs) have also been used to identify and classify images at or above human performance^23^. In the biomedical field, these models have been developed for applications such as cell segmentation^24^, cell classification^25,26^, and tissue segmentation^27^, including our previous work to classify cardiac fibroblasts on the binary scale^28^. Lastly, recent advances in deep learning models have helped to remove human bias from scientific systems. Termed self-supervised learning, models such as BYOL^29^ work to learn similarities and differences between image classes without the need for manually assigned labels. Altogether, these algorithms have revolutionized the field of pattern recognition in biomedicine using features readily obtained from microscopy images.

Interestingly, one method often used to quantify differential expression of cellular features is single-cell RNA sequencing^30^ (scRNA-seq). Analyzing the transcriptome at a single cell level provides much higher resolution data and makes possible the identification of minority populations of cells that would be lost in the noise of bulk experiments, similar to how classifying myofibroblasts on a binary scale loses a diverse population of intermediate phenotypes. This technique has been used to identify small populations of cells in highly heterogeneous environments such as the heart^31,32^ and to characterize in great detail how the cellular composition the heart^33–36^ and lungs^37,38^ change in response to fibrotic disease. However, despite recent advances, scRNA-seq still remains a difficult and costly experiment to conduct, and microscopy-based analyses such as morphological profiling^39–42^ could facilitate faster characterization of heterogeneous samples.

In this work, we report a detailed image-based characterization of cardiac fibroblasts, activated myofibroblasts, and the range of phenotypes between the two. We show that there are significant differences in many simple size and shape features between cells of the two phenotypes, and that these features provide more than enough information to accurately classify each individual cell as either activated or not, as compared to traditional manual classification using α-SMA stress fiber organization. Next, we also use these features to create a model to quantify the position of cells on the continuous spectrum of activation. This model provides more detailed information on the behavior of individual cells and is much more representative of the activation process. Furthermore, we use self-supervised machine learning methods to remove human bias in the continuous classification process. Finally, we demonstrate that this spectrum of activation is not only seen using imaging methods but is strongly correlated to results measured by single-cell RNA sequencing.

## 2. Materials and Methods

### 2.1 Cell Culture

Human cardiac fibroblasts (Lonza, NHCF-A) were cultured on 100 mm tissue culture plastic polystyrene petri dishes in complete DMEM with 10% fetal bovine serum (Corning) and 1% penicillin/streptomycin (Fisher Scientific). Cells were cultured between 4 and 10 passages before passaging onto a #1.5 glass bottom mini petri dish (Idibi) for fluorescent and phase contrast imaging.

### 2.2 Immunostaining and Microscopy

Cells were first fixed in a 2% para-formaldehyde in PBS solution for 10 min, then permeabilized in a 0.2% Triton in 2% para-formaldehyde solution for 3 min. Cells were then blocked in 1% BSA buffer for 1 h on a shaker table. Primary antibody (mouse anti – α-SMA, 3 μg/mL, Abcam) was then added and cells were left at 4 °C overnight. The next day, cells were washed 3 x 5 min with PBS before adding secondary antibody (1:200 Alexafluor-488 goat anti-mouse, Invitrogen) and rhodamine phalloidin (1:100, Invitrogen). Cells were then placed on a shaker table for 1 h at room temperature. Cells were then washed 3 x 5 min, stained with DAPI (1:1000, Invitrogen) for 10 min, and finally washed 2 x 5 min. All imaging was performed on a Nikon Ti2-E eclipse microscope with a 20x objective.

### 2.3 Single-cell RNA sequencing (scRNA-seq) experiment

Cardiac fibroblasts were cultured on petri dishes as previously described. At passage 5, 5 plates of cells were cleaved with 0.25% trypsin solution (Corning). Cells were then spun down into a pellet and resuspended in 1% BSA/PBS at a concentration of 1,000 cells/μL. Cells were then placed on ice for ~45 min before targeting 5,000 cells on the Chromium Controller instrument using the Chromium Next GEM Single Cell 3’ reagent Kit, v3.1 (10x Genomics) according to the manufacturers protocol. Single cell partitioning, cDNA library preparation, and sequencing using a NovaSeq6000 was performed by the University of Texas Genomic Sequencing and Analysis Facility. An Agilent Bioanalyzer (Agilent) and the KAPA SYBR FAST qPCR kit (Roche) were used to determine the quality and concentration of the finished library. Libraries were sequenced to a depth of 50,000 reads per cell.

### 2.4 scRNA-seq Data Processing and Analysis

The raw sequencing data was demultiplexed and aligned using10x Genomics Cell Ranger 6.1.2^30^. The reads were aligned to the Human Reference Genome GRCh38 version 2020-A from 10x Genomics and counted using the Cell Ranger count pipeline. The resulting filtered gene-barcode matrices were then further analyzed using the Seurat package^43^ in R version 4.1.2^44^. To remove low-quality cells, empty droplets, and cell multiplets, cells that had >10% mitochondrial counts, <1500 unique features, or >150,000 total counts were filtered out. The SCTransform function^45^ was then used for normalization to remove the influence of technical variation from downstream analysis and regress out percent of mitochondrial gene mapping. The normalized data was then reduced to the top 50 PCs by PCA and visualized with UMAP^46^. Cells were clustered using Seurat FindClusters function with a resolution of 0.8. Differential gene expression analysis was then performed between top clusters of interest using the Seurat FindMarkers function.

### 2.5 Computation

Cell features were extracted from images using Matlab. Matlab was also used to create the decision-tree, kNN, and SVM models. All other code was written in python. Self-supervised/BYOL model was trained on the Longhorn supercomputer at the Texas Advanced Computing Center. All code is available at the following GitHub repository: https://github.com/ahillsley/cont_classification

### 2.6 Statistical Analysis

For measured cell features, significance was determined using a two-sample, 2 tailed, t-Test assuming equal variances.

## 3. Results

### 3.1 Morphological Profiling of Cardiac Fibroblasts and Myofibroblasts

A dataset was constructed of 1170 individual human cardiac fibroblast cells. Cells were cultured on glass bottom petri dishes between 3 and 10 days before staining and fixation, resulting in a mixed population with fibroblast and myofibroblast phenotypes. While glass has been shown to activate cells similarly to chemical stimuli such as TGF-β1^47^, this activation is far from complete. Many cells remain non-activated and are visually indistinguishable from cells grown on a soft substrate, such as those seen in our previous work^28^. Thus, these samples represent a heterogeneous population. Each cell was imaged through 4 different channels: blue/DAPI to visualize the nucleus, red/F-actin for the cellular cytoskeleton, green/α-SMA for the specific actin isoform associated with myofibroblast activation, and in phase contrast (Figure 1A). Phase contrast was included because any information gained from that channel would be compatible with future live cell experiments. After imaging, individual cells were manually segmented from each image (approx. 3 cells per original image).

**Figure 1:**
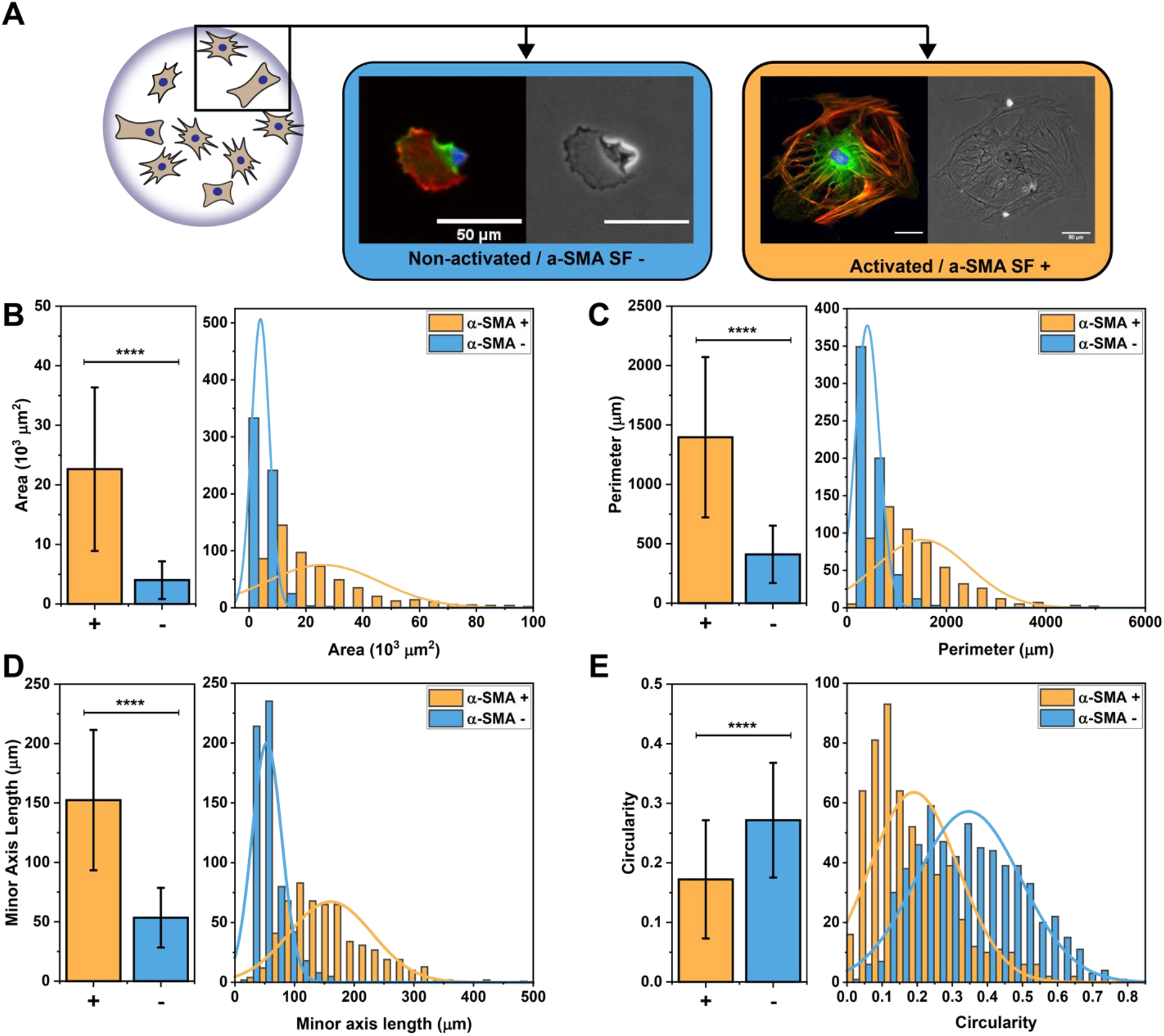
A) Representative 3 channel fluorescent (Red:F-actin, Green:α-SMA, Blue:DAPI) and phase contrast images of myofibroblasts and fibroblasts (scale bar = 50μm) B,C,D,E) Bar plots and histograms showing the averages and distribution of four cell size and shape features: area, perimeter, minor axis length, and circularity, respectively. All of these features were significantly different between the two cell phenotypes. N=566 activated cells and 604 non-activated cells, error bars = standard deviation, **** p < 0.0001.

Initially, each cell was then manually classified as either an α-SMA stress fiber positive activated myofibroblast or as an α-SMA stress fiber negative non-activated fibroblast. Overall, our dataset consisted of 566 activated myofibroblasts and 604 non-activated fibroblasts. This classification was based on the *de novo* appearance of stress fibers in the alpha-smooth muscle actin (green) channel. Cells exhibit a range of α-SMA expression, so while the majority of cells were easily classified, a significant population existed in an intermediate phenotype that required the author’s discretion for classification.

In order to more quantitatively evaluate our cell population, we created a Matlab script to compute 15 different characteristic features for each cell. These calculations were performed on binarized versions of the red/F-actin channel and the blue/DAPI/nuclear channel. The features calculated include area, perimeter, major and minor axis length, circularity, and eccentricity for both the cell and nucleus. Additional features include the cell extent, nuclear/cytosolic ratio, and Minkowski-Bouligand dimension (Table S1).

Significant differences (p < 0.0001) were seen between the two cell phenotypes in 14 out of 15 of the features measured (all except cell extent, Table S1). Cell area was the most significant feature with myofibroblasts on average over five-fold larger than fibroblasts (22,000 μm^2^ vs 4000 μm^2^) (Figure 1B). Other cell size features followed a similar trend, with cell perimeter (Figure 1C, 1397 μm vs 411 μm) and minor axis length (Figure 1D, 242 μm vs 114 μm) being significantly higher for myofibroblasts compared to fibroblasts. The size of the nucleus was also significantly larger in myofibroblasts than in fibroblasts (485 μm^2^vs 189 μm^2^). Another interesting observation was that fibroblasts were more circular than myofibroblasts (Figure 1E, 0.27 vs 0.17).

While constructing this dataset, it was observed that in cells that exhibited clear a-SMA stress fibers, the α-SMA stress fibers appeared to be colocalized with the F-actin cytoskeletal fibers. This phenomenon has been previously observed;^48^ however, to the best of our knowledge, this colocalization has not been explored as a feature in myofibroblast classification models. To further investigate this, we created a python script to quantify the degree of colocalization using Pearson’s correlation coefficient, R_P_ (R_P_ =1 is perfect colocalization, while 0 is random organization). To evaluate the degree of colocalization, we divided the cell into small square tiles, ranging from 4×4 to 64×64 pixels and calculated a RP value for each tile, then averaged all the tiles in the image to determine a value for each cell (Figure S1). Importantly, we masked the cell using the binarized F-actin channel, so as to only consider pixels within the cell. Activated myofibroblasts exhibited a significantly greater R_P_ value than non-activated fibroblasts (0.47 vs 0.11 for 64-pixel tiles) across all tile sizes (Table S1), indicating a greater degree of colocalization between a-SMA and F-actin.

In summary, our dataset of over 1,000 individual cells characterizes and quantifies the morphological differences between activated and non-activated cardiac myofibroblasts cultured on glass. We also demonstrated that there is a statistical difference between morphological features for cells in these two populations. However, when both populations are pooled together, all of our metrics form a single distribution rather than a bimodal one (Figure S2), suggesting that activation is not best characterized by a binary classification, but rather exists on a continuous spectrum.

### 3.2 Machine Learning to Predict Cardiac Fibroblast Activation

We next used these morphological features to develop a quantitative, well-defined method of classifying cells as either activated or non-activated. Towards that goal, we created Reciever Operating Characteristic (ROC) curves and calculated the Area under the Curve (AUC) values to evaluate each of these cell features, and their usefulness for differentiating between activated and non-activated cells (Table 1, Figure S3). Unsurprisingly, cell size parameters, such as area, perimeter, and minor axis length, displayed the greatest ability to differentiate between cell types, with cell area having the highest AUC value of 0.97. Our colocalization metric RP also proved to be able to differentiate with an AUC value of 0.90 for 64×64 pixel tiles, and similar values for all other tile sizes.

**Table 1:**
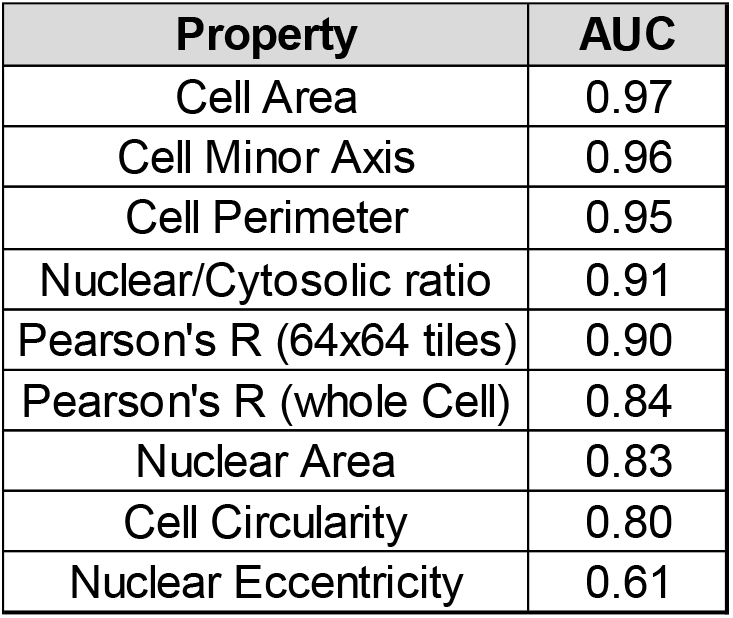
The area under the curve (AUC) of 9 different manually engineered features shows many are good predictors of cell phenotype

Moving forward with cell area as our best feature, we classified cells as activated if they were larger than our we classiiied cells as activated if they were larger than ou cutoff value and as non-activated if they were smaller. After cycling through all measured cell areas, the optimal cutoff value was determined to be approximately 8,000 μm^2^, which alone yields an accuracy of 76% when compared to our manual activation predictions based on the appearance of α-SMA stress fibers. This cell size is similar to that of activated cells cultured on stiff hydrogel substrates^28^. This result was promising, but still too inaccurate to be used to automate the classification of cell activation.

We next used three different machine learning algorithms to increase the prediction accuracy by combining information from multiple cell features. For each of the following models, we decided to move forward only with a select few features that can be derived solely from cell shape (cell area, perimeter, minor axis length, and circularity, Figure 1B-E). This was done to remove the reliance of this technique on cell fixation and staining. One of the largest experimental limitations to studying the dynamics of the myofibroblast transition is the need to fix and stain cells to determine their activation. This is a time-consuming process and results in high sample numbers to observe trends over a time course. Use of any of the following models only requires the determination of cell shape, which can be done through the use of a cytocompatible cell membrane stain or through phase contrast imaging. This opens the possibility of tracking individual live cells through the entire activation process and classifying activation in real time.

The original 1170 single cell images were first randomly split into a training set and test set. Each set was balanced with a relatively even number of activated and non-activated cells. The training set (1,000 cells) was then used to develop the models, while the test set (170 cells / images) was withheld and only used later to evaluate model performance (Table 2).

**Table 2:**
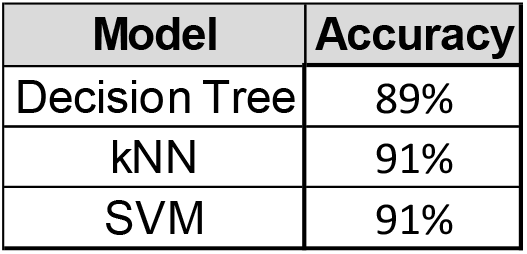
The performance of 3 simple machine learning models predicting the binary label, when provided a short vector of engineered cell features

| Model | Accuracy |
| --- | --- |
| Decision Tree | 89% |
| kNN | 91% |
| SVM | 91% |

The first model we developed was a decision tree (diagrammed in Figure S4). The software JMP was used to empirically determine four different cutoff values that when combined can effectively label the activation of each cell. Adding these three other features in addition to cell area increased the accuracy of our model to 89%. We next used the scikit-learn python package to construct a k-nearest-neighbor (kNN) classifier. This model resulted in a classification accuracy of 91%. Lastly, we also used scikit-learn to construct a Support Vector Machine (SVM) classifier. This model also resulted in a classification accuracy of 91%.

With these models, we have effectively quantified the morphological features of cardiac myofibroblasts and used those features to establish a quantitative process of fibroblast/myofibroblast classification. The developed model matches the α-SMA stress fiber method of cell classification solely from cell size and shape features. However, these cells are still classified on a binary scale, while it has been shown that activation exists on a continuous spectrum. Furthermore, the developed model is compared to the author’s manually assigned labels, which contain inherent bias and some degree of subjectivity.

### 3.3 Classifying cardiac fibroblast activation on a continuous scale

Each cell in our dataset is associated with a 4D vector containing the four shape features previously mentioned (cell area, perimeter, minor axis length, and circularity). Our primary goal is to use this feature vector to create a new system of continuous labels (Figure 2A). To achieve this, we performed principal component analysis (PCA) on the cell feature matrix (Figure 2B). PCA reduces the data, in order to capture the most variance in the minimum number of dimensions. For these manually engineered features, the first principal component (PC 1) accounts for 88% of the total variance. A continuous labeling system was thus created by determining the position of each cell along PC 1 and re-scaling it between 0 and 1,000 (to provide a measure using round numbers).

**Figure 2:**
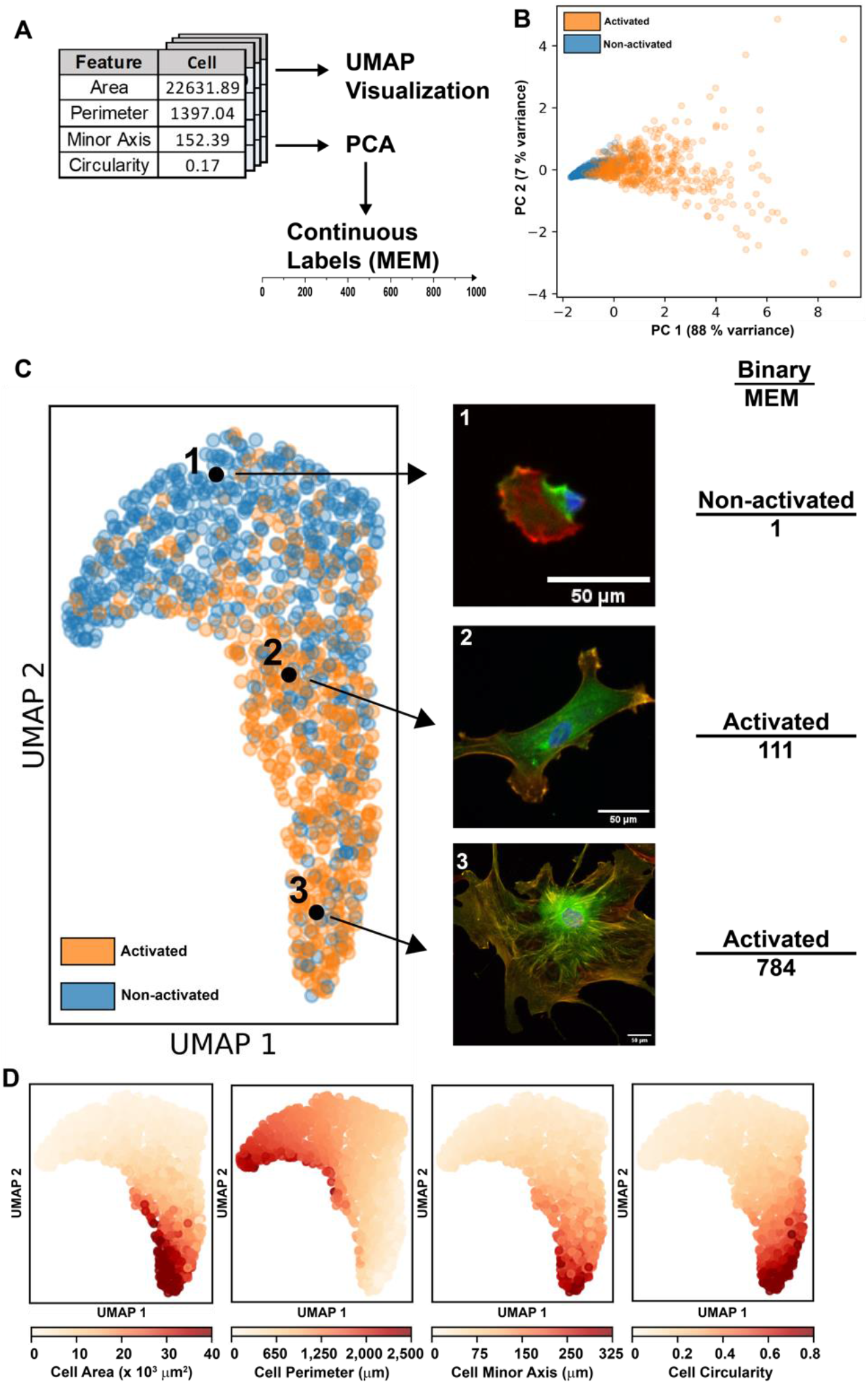
A) Overview of the analytical pipeline. Cell feature vectors were first visualized using UMAP, then reduced using PCA and re-scaled to create a continuous label system. B) 2D PCA reduction of the cell feature vector, PC 1 contained 88% of the variance and was used to create the MEM labels. C) UMAP reduction of the manually engineered features of all 1104 cells, highlighting cells of different activation levels, with both their binary and MEM label. D) Labeling the UMAP reduction by cell features helps to understand how cells are organized in the reduction.

In order to better visualize the distribution of these cells, we first expanded the number of features by including all linear combinations of the four features, then reduced the vector to two dimensions. This was done through the use of Uniform Manifold Approximation and Reduction (UMAP) (Figure 2C). UMAP works by first constructing a high-dimensional representation of the data, then optimizes a lower (2) dimensional graph to be as structurally similar as possible. It is important to note that the orange and blue activation state labels were manually added after the fact for clarity and did not play any role in determining the position of any individual cell. Therefore, this 2D clustering of individual cells is relatively free from external bias; the only existing bias is in the selection of the features themselves and the hyperparameters in the UMAP algorithm. As we can see, based solely on these four size and shape features, the cells are organized in a continuous spectrum roughly along the UMAP 2 axis.

Figure 2C highlights three cells selected from different positions on this activation spectrum. Under the binary classification system, cells 2 and 3 may both be classified as activated myofibroblasts because they both display at least a single α-SMA stress fiber. However, these cells have significant differences that this classification system cannot account for, i.e. cell 3 is significantly larger and has many clearly defined stress fibers compared to fewer, less developed fibers in cell 2. However, the proposed continuous classification system is much better equipped to capture these differences. Cell 3 is scored 748, on the very upper end of the activation scale, while cell 2 is scored 111, closer to the middle of the distribution and much less activated than cell 3. Because the features in the model were specifically chosen to offer the most variation between phenotypes, we will refer to this as the Manually Engineered Model (MEM) and this set of labels as “MEM labels.”

To better visualize how these individual cells are clustered in the UMAP reduction, heat maps were generated by labeling cells based on specific properties shown to vary significantly with activation in section 3.1 (Figure 2D). As expected, cells with a larger cell area are organized towards the bottom of the spectrum, while those properties decrease significantly as UMAP_2 increases. Importantly though, cells are not organized exactly by increasing area, demonstrating that this is not the only property that matters, and the other three features do play a significant role in determining cell position. Interestingly, and in contrast to the other properties, cell circularity is shown to vary significantly along UMAP_1 (Figure 2D). This makes sense given that circularity is not a strong predictor of activation (AUC = 0.8), and activation primarily varies with the UMAP_2 axis.

This continuous classification provides more detail about the activation process and is able to describe intermediate phenotypes. Additionally, the trained MEM model only requires basic cell shape and size features as inputs. This eliminates the need for fixing and staining of specific cell structures to determine cell activation level. However, a significant source of bias still exists in that the MEM model was trained using the author’s definition of activation (i.e., presence of α-SMA stress fibers).

### 3.4 Self-supervised cell classification

In order to remove the authors’ personal biases, we next turned to self-supervised learning, specifically to a method called “Bootstrap Your Own Latent” or BYOL. BYOL works by first duplicating an image and augmenting it so that the new image is similar, but not identical to the original (ie. rotated/flipped/cropped). Both images are then passed through encoders, in this case a ResNet_50, and then projected into a vector representation. A contrastive loss function is then used to minimize the difference between the vector representation of the original image and the augmented image. As a result, the model learns to cluster together the vector representations of similar images, while distancing them from the representations of different images. Because these vector representations are large 1D vectors, we can think of each value as an abstract feature that the model has learned. This vector is then functionally the same as our list of cell features measured in section 3.1; however, in this case we have 2048 abstract features rather than 4 manually measured features for the MEM. We can then once again use UMAP to reduce the dimensionality of this feature vector and visualize how the images are clustered. In addition, we can use PCA to develop a continuous labeling scale. The model created by this pipeline is hereafter referred to as the self-supervised model (SSM, Figure 3A).

**Figure 3:**
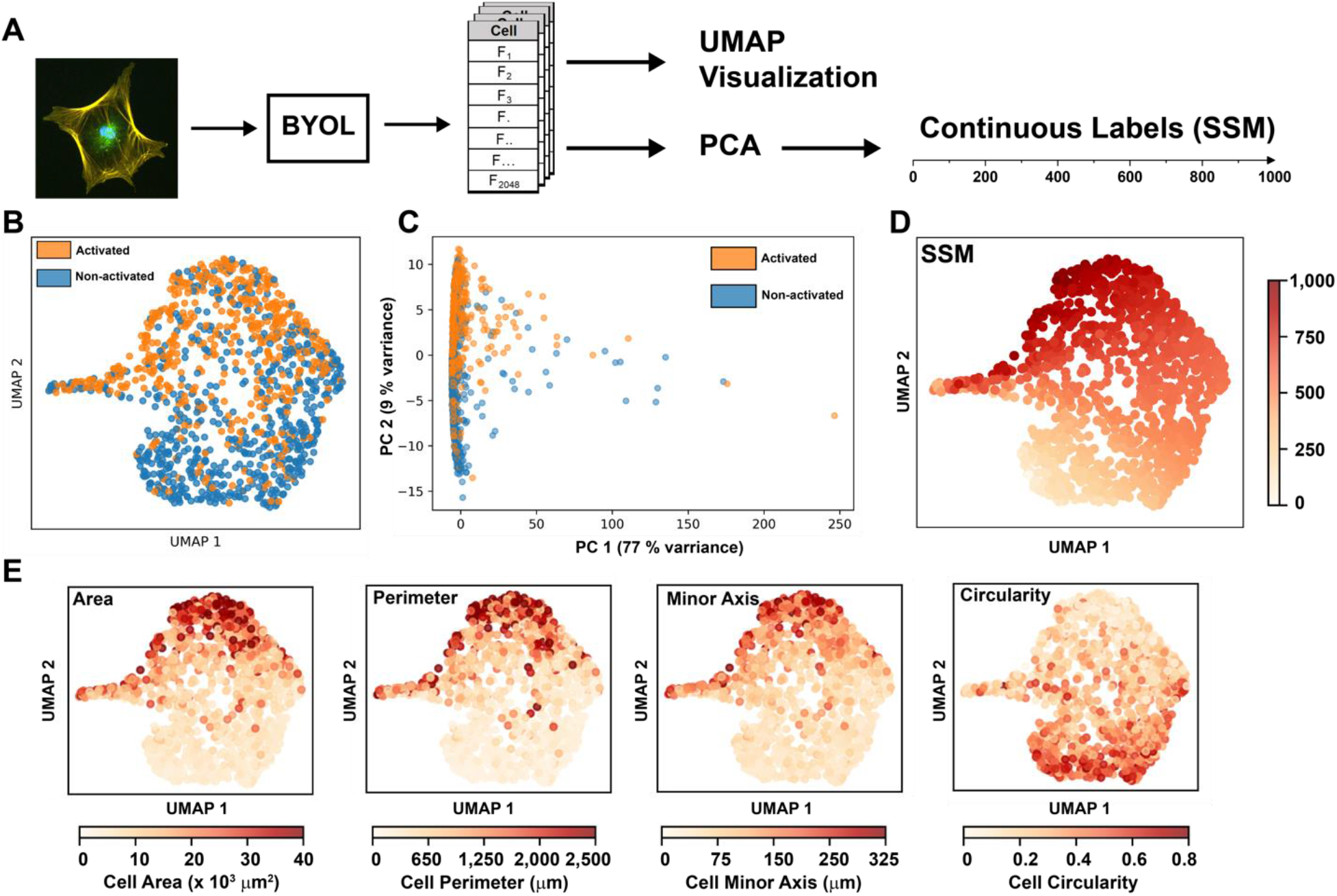
A) Overview of the analytical pipeline. Cell images were first normalized, then used as inputs to train a BYOL model. The new abstract cell features were then visualized with UMAP, and a continuous label system was created using PCA. B) UMAP reduction of the 2048 abstract features learned in the self-supervised model. C) PCA reduction of the abstract cell features. D) Labeling UMAP reduction by SSM labels shows a spectrum of activation. E) Labeling UMAP reduction by cell features shows that this model captures similar patterns to the model in section 3.3.

Importantly, because of how the filters in the model are organized, all images must be the same size; therefore, all of the images were resized to 256 x 256 pixels. As a result, all features learned by the model are independent of cell size characteristics such as area and perimeter, which were the most important predictors of activation from section 3.2. However, even without access to this important information, the model is able to classify cells on a continuous activation spectrum (Figure 3B), very similar to the one seen in Figure 2C. The same method of using PCA to maximize variance (Figure 3C) used in section 3.3 can also be applied to this model, yielding a self-supervised labeling system (SSM). Coloring cells by their SSM labels, shows a clear spectrum of activation increasing in the positive UMAP 2 direction (Figure 3D), which matches the trend seen in the binary labels from Figure 3A. Additionally, by labeling the cells according to the same manually measured features as in Figure 2C and 2D, we can see that the model is arranging the cells in a similar manner to the MEM model (Figure 3E). These SSM labels are now almost completely free from human bias. Interestingly, it appears that the MEM exaggerates differences between highly activated myofibroblasts compared to the SSM, with only 6 cells having a label > 750 in the MEM system, compared to over 250 cells in the SSM system (Figure 4A, 4B, and S5). Additionally, the mean and 95^th^ percentile of the MEM labels is 156 and 483 respectfully, compared to 574 and 889 for the SSM labels. This is likely due to the large tail length of activated myofibroblast size at the top end of the distribution of all cells, which is the most important feature of the MEM, while unimportant to the SSM.

**Figure 4:**
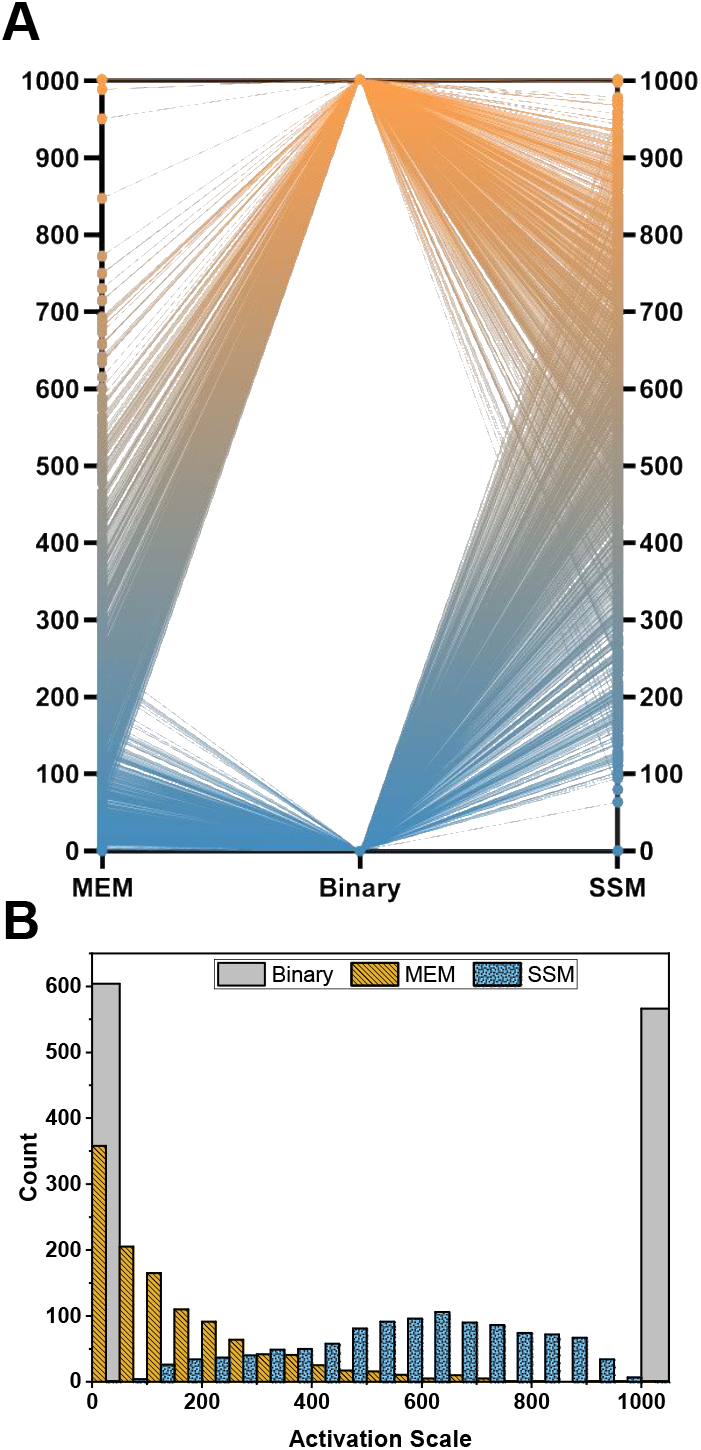
A) A visualization of all the labeling systems shows the increase in resolution gained by the MEM or SSM system over the binary system. B) Histograms showing the distribution of labels for each labeling system.

This SSM labeling convention removes almost all human bias in the classification of these cell phenotypes and clearly demonstrates that based on the fluorescent images, activation is a spectrum and that no clear line exists between an activated myofibroblast and a non-activated fibroblast.

### 3.5 scRNA-seq supports a continuous spectrum of fibroblast activation

In order to support the results of the image-based analyses done in the previous section, we used a complementary experimental technique: single-cell RNA sequencing. Importantly, cells were grown under the same conditions for this experiment as in the prior imaging experiments. The raw sequencing results indicated successful sequencing of 4,062 cells with an average sequencing depth of 77,500 reads per cell and a median of 5,684 genes per cell. After removal of low-quality reads, 3,531 cells remained in the analysis. This experiment resulted in a matrix describing the differing expression levels of over 20,000 specific genes for each individual cell. For each cell, this expression profile can be viewed as a 1 x 20,000 genetic feature vector, similar to the 1 x 2048 abstract feature vector from the SSM or the 1 x 4 cell shape feature vector from the MEM. Using similar dimensionality techniques as in Figures 2 and 3, this genetic feature vector can be reduced to two dimensions for visualization (Figure 5A). As can be seen, most cells belong to a large single cluster oriented along the UMAP 1 direction.

**Figure 5:**
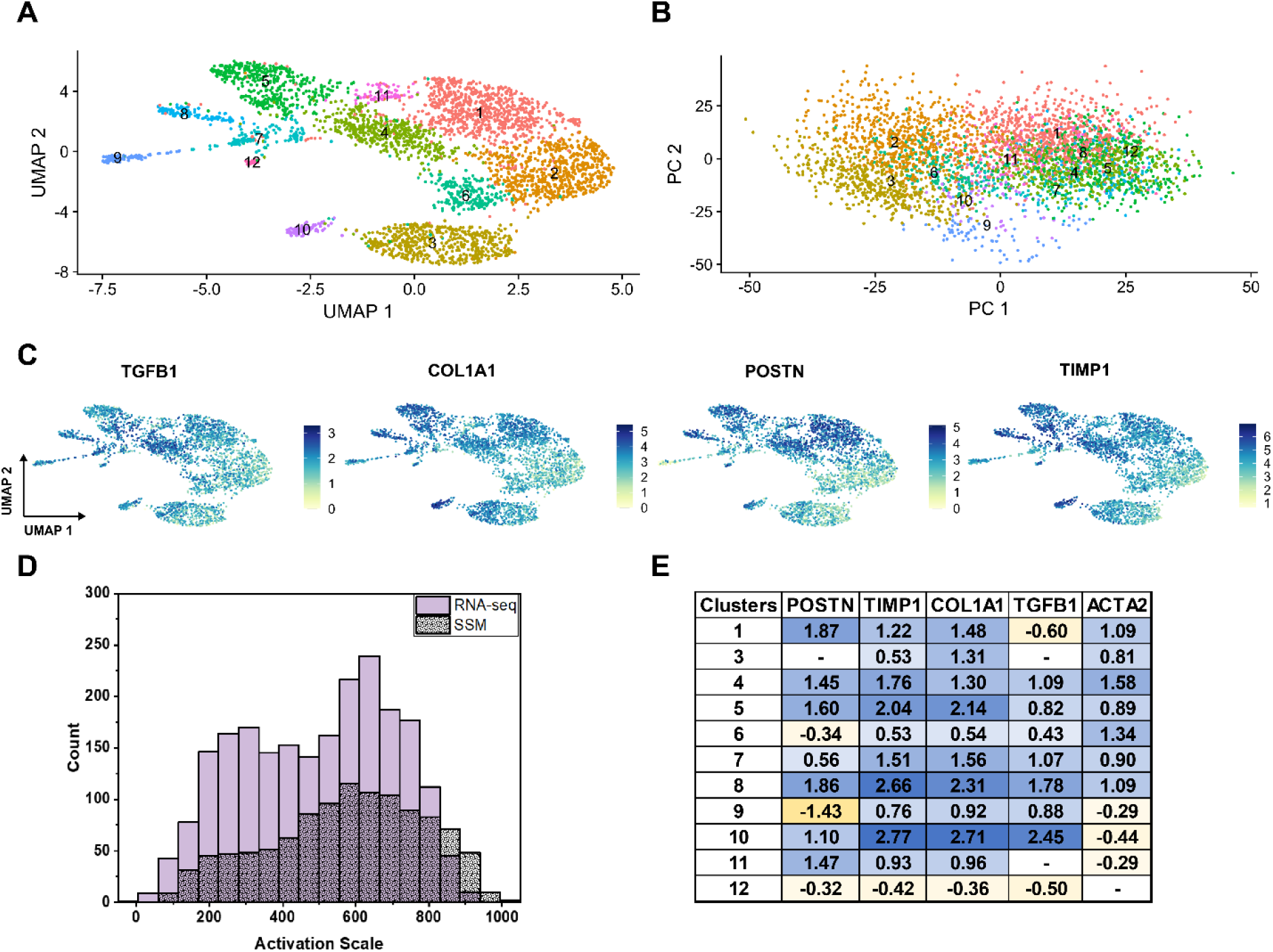
A) UMAP reduction of the transcriptomic feature vector for each cell. Clusters were identified by the Suerat software. B) PCA reduction of the transcriptomic features; individual cells are colored according to their cluster number from (A). C) Labeling each cell by the expression level of four myofibroblast associated genes (TGFB1, COL1A1, POSTN, and TIMP1) shows a consistent spectrum of activation. D) Using the PCA reduction, another continuous label system was created. The distribution of cells is similar to that seen from the SSM model. E) Log2(FC) values of myofibroblast associated genes show that clusters 1, 4, 5, 7, 8, and 10 are highly activated compared to cluster 2.

PCA was also performed on this dataset (Figure 5B) to determine the genes that have the most variance across all cells. A list of genes that compose PC 1 and PC 2 can be found in Table S3, but a few highly differentially expressed genes of interest are: TIMP1^49^, POSTN^12^, TGFB1^50^, and COL1A1^51^, all of which are positively correlated to myofibroblast activation. TIMP1 and POSTN play significant roles in ECM remodeling, the TGFB1 signaling pathway is responsible for myofibroblast activation, and COL1A1 is an important ECM component secreted by myofibroblasts. Coloring the UMAP plot according to the expression of each of these genes (Figure 5C) illustrates a clear increase in expression of each in the negative UMAP 1 direction. This suggests that the genetic distribution of cells is similar to that seen through our image analysis, with activated myofibroblasts on the left-hand side (cluster 5), fibroblasts on the right-hand side (clusters 2) and a range of cells in-between the two extremes. Interestingly, GAPDH was also reported as having significant variance in expression levels, which has also been previously reported^52^. This has significant effects for future work, especially where glass is used as a culture substrate, because GAPDH is often assumed to have consistent expression and is often used as a housekeeping gene for qRT-PCR experiments.

Following a similar pipeline as in sections 3.3 and 3.4, we re-scaled the PC 1 axis from 0 – 1,000 to create another continuous scale of activation (Figure 5D). This scale closely matches the SSM label system, with a mean activation of 573 and a 95^th^ percentile of 889 for the SSM labels, compared to a mean of 501 and a 95^th^ percentile of 795 for the RNA-seq labels. Further both histograms have a mode activation around 600. This verifies that we are capturing the same trends through both imaging and transcriptomic features.

The clusters shown in Figure 5A were automatically generated using the Seurat software, and the differential expression of specific genes between clusters can be quantified by comparing their average log2FC (FC, fold change) values. For example, a log2FC value of 1 corresponds to a 2 fold increase in expression. We compared the log2FC values for the 4 genes of interest and ACTA2 (α-SMA) between each cluster (Figure 5E). Cluster 2 was used as a non-activated baseline, because it displayed the lowest average expression of each of these genes. Clusters 1, 4, 5, 7, 8, and 10 all displayed significantly higher log2FC values than cluster 2, and therefore are assumed to be activated myofibroblasts. Cluster 3, accounting for nearly 15% of the cells, did not show significant differential expression in two of the four selected markers compared to cluster 2, the non-activated baseline. Furthermore, cluster 3 showed small increases in expression of TIMP1 and COL1A1 relative to the clusters identified as activated myofibroblasts, indicating these cells could be in an early stage of activation, further emphasizing the spectrum of activation states. Together, these clusters account for 55.9% of the total number of cells (Table S4). This result is in good agreement with the image dataset collected in section 3.1 which contained 48.4% activated myofibroblasts, identified by a-SMA stress fiber organization. Together with the imaging results, this transcriptomic data supports a continuous spectrum of activation from fibroblast to myofibroblast.

## 4. Discussion and Conclusion

We have used both imaging and transcriptomic techniques to quantify the spectrum of activation from cardiac fibroblast to activated myofibroblast. Features derived from cell images were then used to develop models that are able to predict cell activation at 93.4% accuracy, as compared to manual labels. Importantly, these features are only related to cell shape and size, meaning that it may be possible to accurately determine activation state without the need to fix and stain for specific cellular structures. This provides the possibility of tracking individual live cells at multiple time points throughout the activation process. Next, these features were used to propose a new continuous labeling system (MEM) that provides much higher resolution information about the cells in a given system than the standard binary classification and is more representative of the results seen from transcriptomic analysis.

An interesting observation is that the MEM labels are significantly skewed, when compared to the other labeling systems, with an average activation of 150 / 1,000. This is caused by the large role cell area plays in the PCA reduction. As seen in Figure 1B, the distribution of cell area has a very large tail with a few cells being up to 10x larger than the average. This wide distribution is reflected in the PCA reduction (Figure 2B). Therefore, the scaling of MEM labels from 0 to 1,000 results in only a few number of cells > 800, and the average cell at ~150 / 1,000. Recognizing this fact, this model still achieves its goals of providing more information about the system than the binary labels.

Next, we used self-supervised machine learning techniques to remove the reliance on human cell classification and feature engineering from this system. The resulting classification system (SSM) is relatively simple to implement and provides a way to standardize results across researchers. It is important to note that in our SSM while cell size features are not used, it is very difficult to assign meaning to the features that are used. The vector representation is highly abstracted and has little relation to traditional features used to characterize cell images. Additionally, while the SSM significantly reduces human bias in the system, it does not completely eliminate all bias. Bias still exists in the choice of images used to train the model and in the choice of UMAP and model hyperparameters. This can be addressed in the future by increasing the number of training images.

Lastly, we used single-cell RNA sequencing to support our imaging-based conclusions with transcriptomic data. A dimensionality reduction of over 20,000 gene expression profiles showed cells organized in a continuous spectrum. Genes associated with myofibroblast activation varied significantly across all cells measured, and displayed a clear spectrum, with roughly 56% of the cells upregulating myofibroblast associated genes.

In summary this work provides the following advances: 1) a continuous scale of activation that is more representative of the activation spectrum, 2) Development of simple methods to classify cells on this continuous scale (MEM), or on the binary scale (SVM or kNN models) without the need for fixation, 3) the reduction of human bias in the classification process (SSM), and 4) verification of trends seen in imaging with transcriptomic data. This work also provides a general strategy that could easily be applied to many different applications. Researchers wishing to apply this work simply need to create a dataset of individual cell images. Next, a feature vector can be created for each cell, either through our scripts provided on Github, or custom pipelines for dataset specific features. Lastly, PCA and re-scaling to a continuous label system can be done through almost any language (Python, Matlab, etc.).

## Acknowledgments

This research was supported by the Burroughs Wellcome Fund (CASI-1015895, A.M.R) and by the National Institutes of Health (R35GM138193, A.M.R, and 1R21ES032124-01 awarded to L.M.C.). This material is based upon work supported by the National Science Foundation Graduate Research Fellowship Program awarded to S.M.E. under Grant No. DGE 2137420. Any opinions, findings, and conclusions or recommendations expressed in this material are those of the author(s) and do not necessarily reflect the views of the National Science Foundation. The authors acknowledge the Texas Advanced Computing Center (TACC) at The University of Texas at Austin for providing high performance computing resources that have contributed to the research results reported within this paper. URL: http://www.tacc.utexas.edu. We also thank the Genomic Sequencing and Analysis Facility at UT Austin, Center for Biomedical Research Support, RRID#: SCR_021713, for help with scRNA-sequencing.

## Author Contributions

A.H and A.M.R designed the research; A.H performed all cell culture, imaging, and led the writing of the manuscript; A.H. and M.S. extracted cell features and built all classification models; S.M.E and L.M.C led scRNA-seq data analysis; A.H, M.S., S.M.E. and A.M.R analyzed data and edited the manuscript.

## Data and Software Availability

All code can be found at the following GitHub page: https://github.com/ahillsley/cont_classification. All training images are available at the Texas Data repository at https://dataverse.tdl.org/dataverse/rosalesche. The complete differential gene analysis between cluster 2 (non-activated fibroblasts) and all other clusters can be found in Tables S5-S15 in the supplemental excel worksheet.

