## Supplemental Information for "A Strategy to Quantify Myofibroblast Activation on a Continuous Spectrum"

### Quantitative analysis of myofibroblast activation and classification on a continuous spectrum

| | Activated | | Non-Activated | |
|---|---|---|---|---|
| **Feature** | **avg** | **error** | **avg** | **error** |
| Cell Area | 22631.89 | 13733.01 | 4002.48 | 3166.87 |
| Cell Perimeter | 1397.04 | 674.56 | 410.99 | 241.26 |
| Cell Major Axis Length | 242.41 | 75.55 | 113.70 | 56.03 |
| Cell Minor Axis Length | 152.39 | 59.06 | 53.46 | 25.08 |
| Cell Circularity | 0.17 | 0.10 | 0.27 | 0.10 |
| Cell Eccentricity | 0.71 | 0.17 | 0.82 | 0.14 |
| Cell Extent | 0.46 | 0.13 | 0.46 | 0.13 |
| Nucleus Area | 485.79 | 319.75 | 189.92 | 96.03 |
| Nucleus Perimeter | 86.94 | 39.45 | 53.32 | 15.27 |
| Nucleus Major Axis Length | 29.14 | 12.15 | 19.27 | 5.12 |
| Nucleus Minor Axis Length | 19.21 | 8.48 | 12.09 | 3.42 |
| Nucleus Circularity | 0.69 | 0.24 | 0.80 | 0.13 |
| Nucleus Eccentricity | 0.68 | 0.21 | 0.75 | 0.12 |
| Nuc Cyt Ratio | 0.02 | 0.01 | 0.04 | 0.01 |
| Minkowski-Bouligand dimension | 1.16 | 0.08 | 1.09 | 0.09 |
| Mean Pearson's r whole | 0.24 | 0.29 | -0.14 | 0.25 |
| Mean Pearson's r 64x64 | 0.47 | 0.20 | 0.11 | 0.20 |
| Mean Pearson's r 32x32 | 0.49 | 0.18 | 0.19 | 0.15 |
| Mean Pearson's r 16x16 | 0.48 | 0.18 | 0.20 | 0.13 |
| Mean Pearson's r 8x8 | 0.44 | 0.17 | 0.18 | 0.11 |
| Mean Pearson's r 4x4 | 0.37 | 0.16 | 0.13 | 0.09 |

**Table S1**. Average value and standard deviation of all measured cell features. Units are in μm for perimeters and axis lengths, $\mu m^2$ for areas, and unitless for all other features.

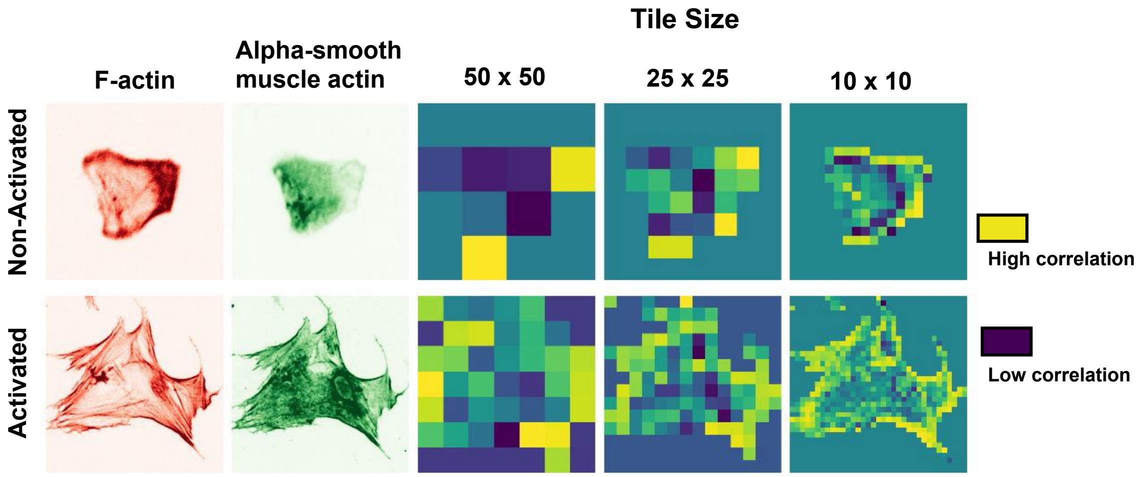


**Figure S1**. Colocalization of F-actin and alpha-SMA was quantified using the Pearson's correlation coefficient ($R_P$). In Python, the image was first broken into a series of small tiles (Tile sizes of 50 x 50 , 25 x 25, and 10 x 10 pixels shown). $R_P$ was then calculated for each tile (yellow – high colocalization, purple – low colocalization). All cell containing tiles were then averaged to calculate an average colocalization for each cell. On average, activated cells had a significantly higher degree of colocalization across all tile sizes.

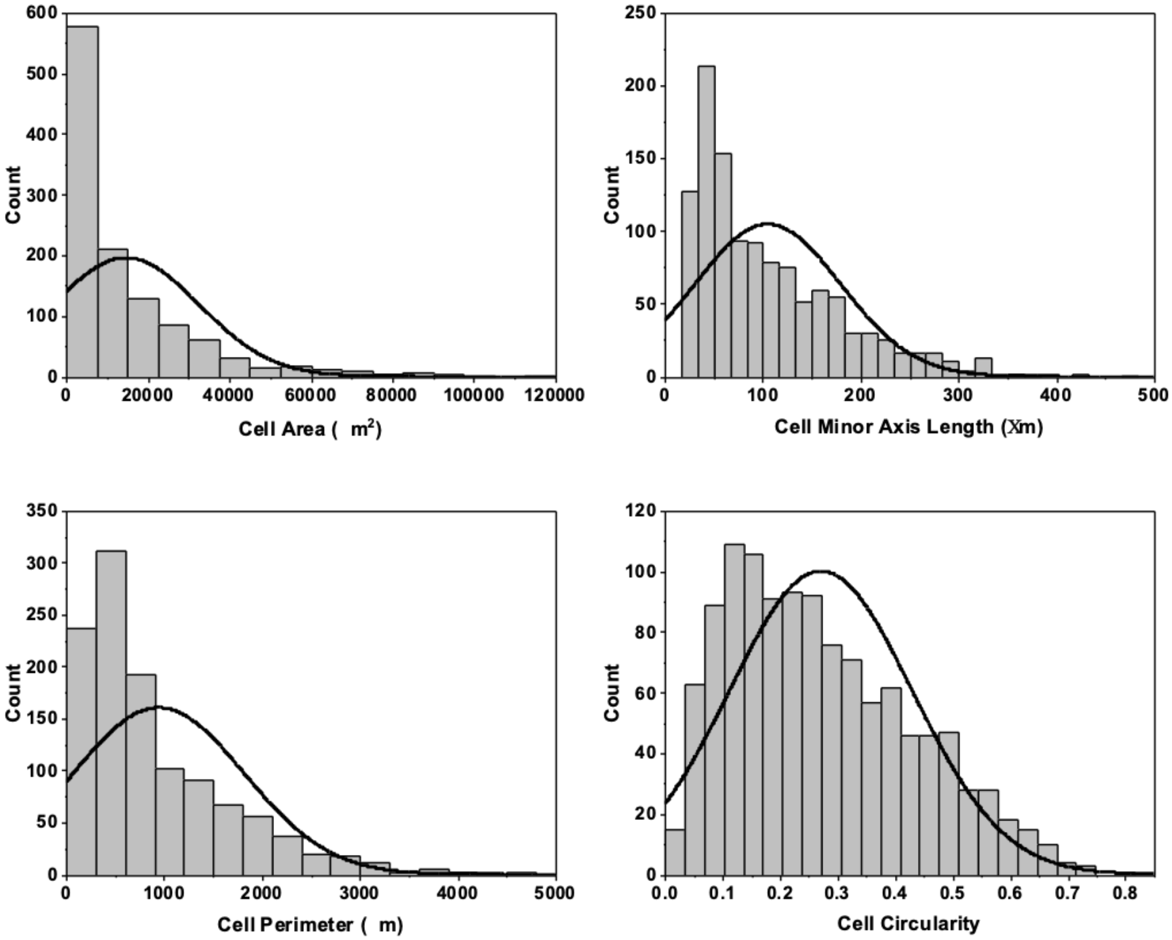


**Figure S2**. Histograms of four measured cell features for all 1170 cells. Without manually separating activated myofibroblasts from non-activated fibroblasts, the cells appear to be a part of a single distribution.

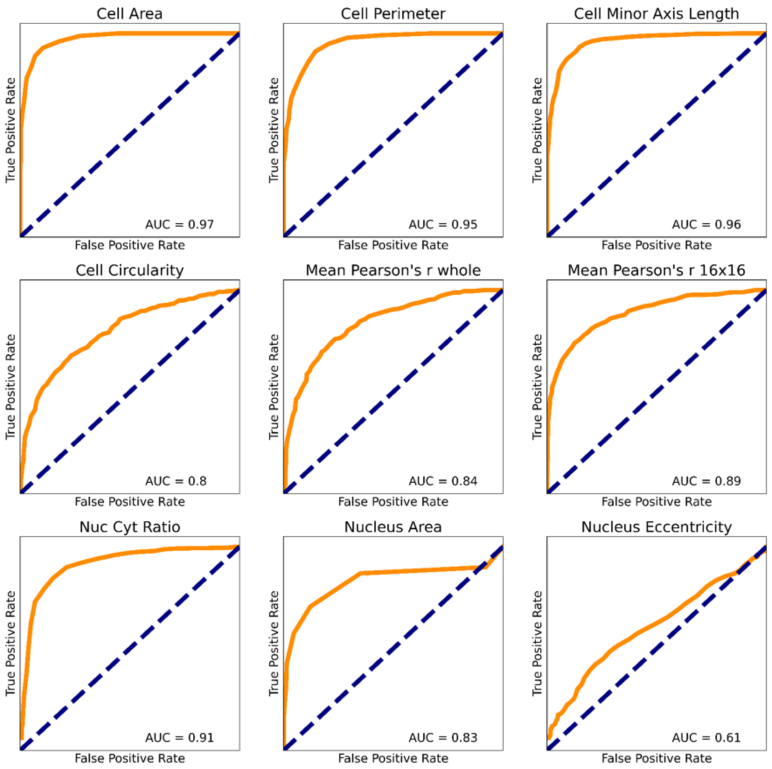


| **Property** | **AUC** |
|---|---|
| Cell Area | 0.97 |
| Cell Minor | 0.96 |
| Cell Perimeter | 0.95 |
| Cell Major | 0.92 |
| Nuclear/Cytosolic ratio | 0.91 |
| Pearson's R 64x64 | 0.90 |
| Pearson's R 8x8 | 0.89 |
| Pearson's R 32x32 | 0.89 |
| Pearson's R 4x4 | 0.89 |
| Pearson's R 16x16 | 0.89 |
| Pearson's R whole | 0.84 |
| Nuclear Area | 0.83 |
| Nuclear Major | 0.82 |
| Nuclear Minor | 0.82 |
| Nuclear Perimeter | 0.81 |
| Cell Circularity | 0.80 |
| Minkowski–Bouligand dimension | 0.72 |
| Cell Eccentricity | 0.69 |
| Nuclear Circularity | 0.66 |
| Nuclear Eccentricity | 0.61 |
| Cell Extent | 0.51 |

**Figure S3**. ROC curves for nine measured cell features. AUC values were calculated from ROC curves for all 21 features and are shown in the table.

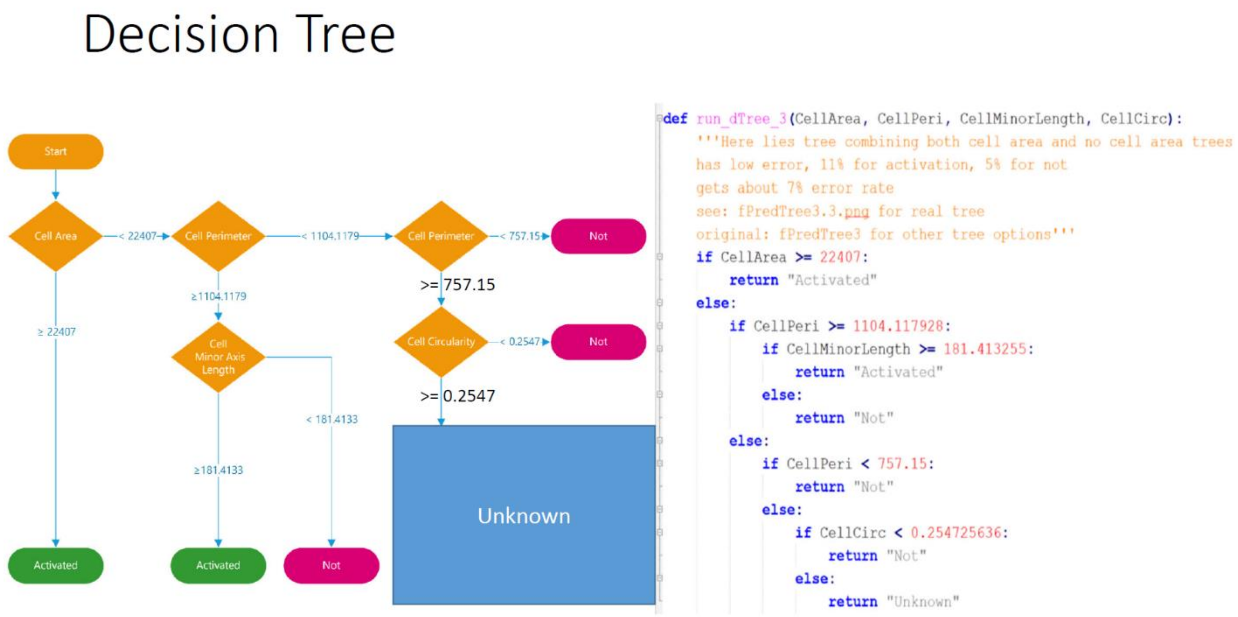


**Figure S4**. Illustration of the decision tree model and the "if" statements used in its code.

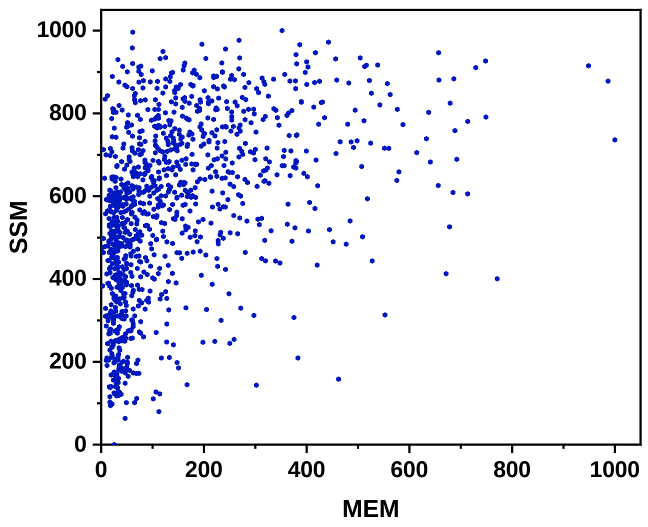


**Figure S5**. Comparison of the SSM and MEM label systems. SSM provides a relatively even distribution of labels between 0 and 1,000, while the MEM labels are clustered at the lower end of the spectrum, with very few cells labeled > 800.

| PC 1 | | | PC2 | |
|---|---|---|---|---|
| **Up-Regulated** | **Down-Regulated** | | **Up-Regulated** | **Down-Regulated** |
| MT2A | MALAT1 | | PTX3 | S100A16 |
| KRT18 | GAS5 | | ANKRD1 | S100A6 |
| TIMP1 | S100A16 | | CCN2 | AKR1C1 |
| UACA | FGF2 | | ADAMTS1 | AKR1C3 |
| POSTN | RALA | | FGF2 | S100A4 |
| CRYAB | MFAP5 | | MALAT1 | MFAP5 |
| IGFBP5 | DDIT3 | | PPME1 | TMEM158 |
| PLAT | THBS1 | | TNFRSF11B | GAPDH |
| TFPI2 | VIPR1 | | THBS1 | S100A10 |
| PRSS23 | NIBAN1 | | INHBA | NRP2 |
| LGALS1 | ZFAS1 | | TPM1 | TIMP1 |
| MYL9 | SH3BGR | | KRT18 | FTL |
| SPARC | NRP2 | | SERPINE1 | AKR1C2 |
| GAPDH | PTX3 | | PCDH10 | VIM |
| HSPB6 | TGFB2 | | CALD1 | FTH1 |
| SH3BGRL3 | HSPA5 | | COL8A1 | RARRES2 |
| TPM2 | AKR1C1 | | MMP1 | CTHRC1 |
| SCUBE3 | GDF15 | | FST | PDLIM4 |
| TGFB1 | SLC3A2 | | PCDH17 | CLEC2B |
| MGP | SYT1 | | UACA | AKR1B1 |
| COL1A1 | NEAT1 | | CDC42EP3 | LGALS3 |
| VIM | GADD45A | | ANK3 | CTSK |
| PRSS3 | C5orf46 | | UGCG | MEG3 |
| CD59 | NNMT | | RGS4 | MMP3 |
| TMSB10 | EFEMP1 | | SYNE1 | SPON2 |
| THY1 | EPAS1 | | DSP | CFH |
| TUBB3 | IFI16 | | RND3 | TIMP3 |
| SERF2 | TRIB3 | | STAT1 | S100A13 |
| ACTB | AKR1C3 | | DDIT3 | SPP1 |
| IGFBP7 | PCDH10 | | POSTN | KCTD12 |

**Table S3**. List of genes significantly up and down regulated along PC 1 and PC 2 axes of the scRNA-seq analysis. Upregulated genes become more highly expressed in the positive direction of PC 1 and PC 2, while the opposite is true for downregulated genes. These genes contribute the most to the variance seen between all measured cells.

| Cluster | # of Cells | % Total |
|---|---|---|
| 1 | 776 | 22.0% |
| 2 | 648 | 18.4% |
| 3 | 517 | 14.6% |
| 4 | 458 | 13.0% |
| 5 | 431 | 12.2% |
| 6 | 219 | 6.2% |
| 7 | 121 | 3.4% |
| 8 | 112 | 3.2% |
| 9 | 87 | 2.5% |
| 10 | 75 | 2.1% |
| 11 | 67 | 1.9% |
| 12 | 20 | 0.6% |
| Total | 3531 | |

**Table S4.** The total number of cells in each cluster of Figure 4A. Clusters were generated automatically using the Seurat software.
